# LTR_retriever: a highly accurate and sensitive program for identification of LTR retrotransposons

**DOI:** 10.1101/137141

**Authors:** Shujun Ou, Ning Jiang

**Affiliations:** Department of Horticulture, Michigan State University, East Lansing, MI, 48824, USA

**Keywords:** LTR retrotransposon, LTR_retriever, transposable element, genome annotation, evolution

## Abstract

Long terminal-repeat retrotransposons (LTR-RTs) are prevalent in plant genomes. Identification of LTR-RTs is critical for achieving high-quality gene annotation. Based on the well-conserved structure, multiple programs were developed for *de novo* identification of LTR-RTs; however, these programs are associated with low specificity and high false discovery rate (FDR). Here we report LTR_retriever, a multithreading empowered Perl program that identifies LTR-RTs and generates high-quality LTR libraries from genomic sequences. LTR_retriever demonstrated significant improvements by achieving high levels of sensitivity (91.8%), specificity (94.7%), accuracy (94.3%), and precision (90.6%) in model plants. LTR_retriever is also compatible with long sequencing reads. With 40k self-corrected PacBio reads equivalent to 4.5X genome coverage in Arabidopsis, the constructed LTR library showed excellent sensitivity and specificity. In addition to canonical LTR-RTs with 5'-TG..CA-3' termini, LTR_retriever also identifies non-canonical LTR-RTs (non-TGCA), which have been largely ignored in genome-wide studies. We identified seven types of non-canonical LTRs from 42 out of 50 plant genomes. The majority of non-canonical LTRs are *Copia* elements, with which the LTR is four times shorter than that of other *Copia* elements, which may be a result of their target specificity. Strikingly, non-TGCA *Copia* elements are often located in genic regions and preferentially insert nearby or within genes, indicating their impact on the evolution of genes and potential as mutagenesis tools.

## INTRODUCTION

Transposable elements (TEs) are ubiquitous interspersed repeats in most sequenced eukaryote genomes (Wessler 2006). According to their transposition schemes, TEs are categorized into two classes. Class I TEs (retrotransposons) use RNA intermediates with a “copy and paste” transposition mechanism (Kumar and Bennetzen 1999; Wicker, et al. 2007). Class II TEs (DNA transposons) use DNA intermediates with a “cut and paste” mechanism (Feschotte and Pritham 2007; Wicker, et al. 2007). Depending on the presence of long terminal repeats (LTRs), Class I TEs are further classified as LTR retrotransposons (LTR-RTs) and non-LTR retrotransposons, including short interspersed transposable elements (SINEs) and long interspersed transposable elements (LINEs) (Han 2010). For simplicity, TEs other than LTR-RT, including both non-LTR retrotransposons and DNA transposons, are called non-LTR in this study. In plants, LTR-RTs contribute significantly to genome size expansion due to their high copy number and large size (Rensing, et al. 2008; Schnable, et al. 2009; Nystedt, et al. 2013; Ming, et al. 2015). For example, retrotransposons contribute to -approximately 75% to the size of the maize (*Zea mays*) genome (Schnable, et al. 2009). In *Oryza australiensis*, a wild relative of rice (*O. sativa*), the amplification of three families of LTR retrotransposons is attributed to the genome size doubling within the last 3 million years (MY) (Piegu, et al. 2006). The amplification and elimination of LTR-RTs has shaped genome landscapes (Ammiraju, et al. 2007; Ammiraju, et al. 2010), thereby affecting the expression of adjacent genes (Hollister and Gaut 2009; Hollister, et al. 2011; vonHoldt, et al. 2012; Makarevitch, et al. 2015).

An intact LTR-RT carries an LTR at both termini (**Fig 1A**). The LTR regions usually span 85-5000 base pairs (bp) with intra-element sequence identity ≥ 85%. In plants, LTRs are typically flanked by 2 bp palindromic motifs (**Fig 1A**), commonly 5'-TG..CA-3' (Zhao, et al. 2016) with some rare exceptions. For instance, the first active TE detected in rice, the *Tos17* LTR element has a 5’-TG…GA-3’ motif (Hirochika, et al. 1996). The sequence between the 5' and 3' LTR is defined as the internal region and usually ranges from 1,000-15,000 bp (**Supplementary Fig S1**). To confer transposition activities, the internal region of most autonomous LTR elements should contain a primer binding site (PBS), a polypurine tract (PPT), a *gag* gene (i.e., encoding structural proteins for reverse transcription), and a *pol* gene (i.e., functioning as protease, reverse transcriptase, and integrase) (Havecker, et al. 2004). Depending on the order of protein domains in the *pol* gene, intact LTR-RTs can be further categorized into two families called *Gypsy* and *Copia* (Kumar and Bennetzen 1999). If the internal region does not contain any open reading frames (ORFs), e.g., reverse transcriptase genes, the belonging LTR-RT is unable to transpose independently, and it relies on the transposition-related proteins from other autonomous LTR-RTs (Havecker, et al. 2004; Jiang 2016). There are two groups of non-coding LTR-RTs: terminal-repeat retrotransposon in miniature (TRIM) (Havecker, et al. 2004; Gao, et al. 2012) and large retrotransposon derivatives (LARD) (Havecker, et al. 2004). These non-coding LTR-RTs are distinguished by their average length: TRIMs are < 1 kb and LARDs are 5.5-9kb (Havecker, et al. 2004; Jiang 2016).

**Fig 1.**
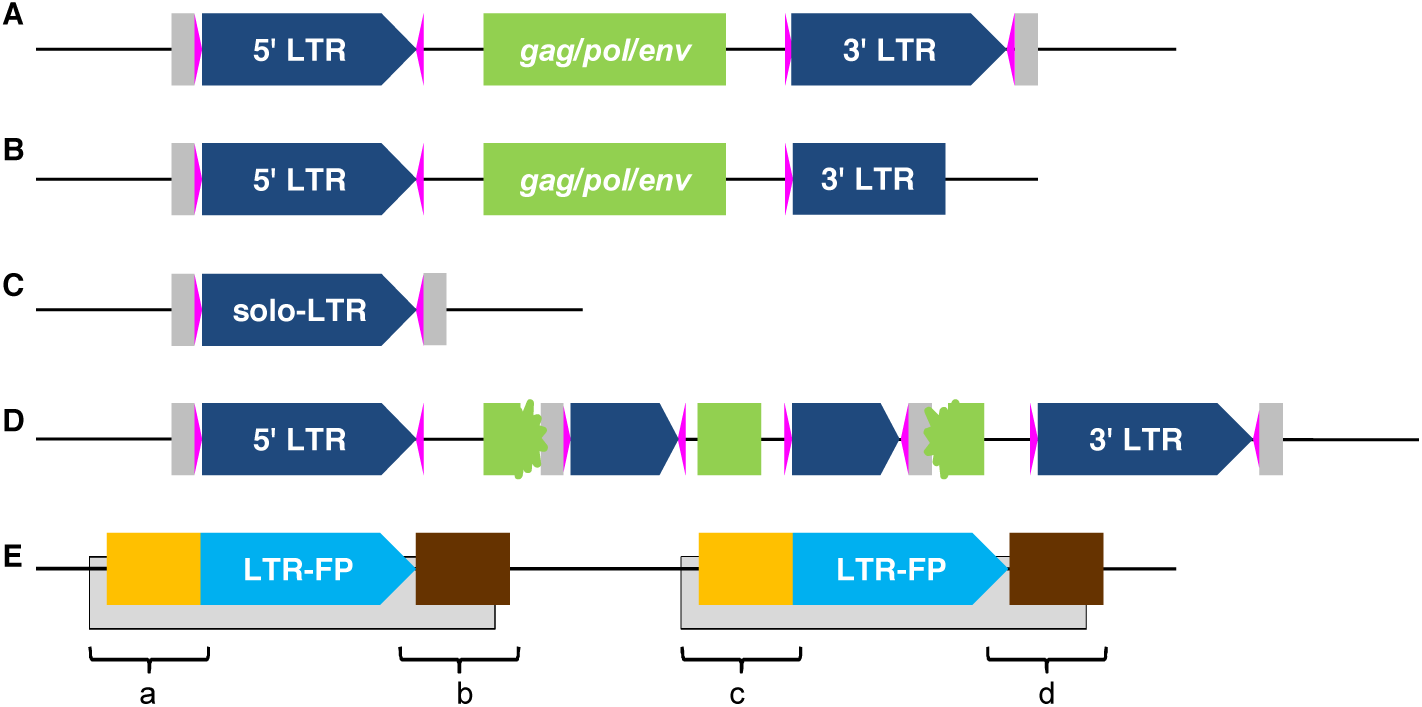
The structure of LTR retrotransposons (LTR-RT), their derivatives, and false positives. (**A**)The structure of an intact LTR-RT with long terminal repeat (LTR) (navy pentagons), a pair of di-nucleotide palindromic motifs flanking each LTR (magenta triangles), the internal region including protein coding sequences for *gag*, *pol*, and *env* (green boxes), and 5 bp target site duplication (TSD) flanking the element (gray boxes). (**B**) A truncated LTR-RT with missing structural components. (**C**) A solo-LTR. (**D**) A nested LTR-RT with another LTR-RT inserted into its coding region. (**E**) A false LTR-RT detected due to two adjacent non-LTR repeats (gray boxes). The counterfeit also features with a direct repeat (blue pentagons) but usually has extended sequence similarity on one or both sides of the LTR (orange and brown boxes). Regions a-d are extracted and analyzed by LTR_retriever.

The insertion of an LTR-RT is accompanied by the duplication of a small piece of sequence immediately flanking the element, which is called target site duplication (TSD, 4-6 bp in length) (**Fig 1A**). There are many mechanisms that can introduce mutations to a newly transposed LTR-RT. Due to the sequence similarity between the long direct repeat of an LTR-RT, intra-element recombination can occur, leading to the elimination of the internal region and the formation of a solo-LTR (**Fig 1C**). The number of solo LTRs indicate the frequency and efficiency of LTR removal in a genome (Tian, et al. 2009). New LTR-RT insertions can be silenced by methylation and chromatin modification as a genomic mechanism to suppress expression (Fedoroff 2012; vonHoldt, et al. 2012). Silenced elements have less selection constraint and accumulate more mutations including deletions, resulting in truncated LTR-RTs (**Fig 1B**). Truncated LTR-RT could also be the product of illegitimate recombination which generates deletions and translocations (Tian, et al. 2009; Zhao, et al. 2016). LTR-RTs often insert into other LTR-RTs, generating nested LTR-RTs (**Fig 1D**) (SanMiguel, et al. 1998; Tian, et al. 2009; Levy, et al. 2010). Given these mutation mechanisms, intact elements only contribute a small fraction of all LTR-RT related sequences in a genome. If the required structural components are altered, i.e., mutated, truncated, and nest-inserted by other TEs (**Fig 1**), the LTR element becomes non-autonomous and is difficult to identify using structural information.

Although the structure of LTR-RT is conserved among species, their nucleotide sequences are not conserved except among closely related species. Particularly, substantial sequence diversity is observed within the long terminal repeat region. Therefore, LTR-RTs are usually not identified based on sequence homology. Due to the lack of nucleotide sequence similarity among species, constructing a species-specific LTR library (i.e., exemplars) is essential for identification of all LTR-RT related sequences in a newly sequenced genome.

Computational identification of LTR-RTs based on structural features has been implemented multiple times. Such methods are usually used jointly to maximize power in genome annotation projects. However, inconsistent results are often obtained from these tools (Hoen, et al. 2015), which could be due to the differences in defining the LTR structure in the program and the different implementation of these methods. LTR_STRUC was one of the earliest developments of genome-wide LTR identification programs (McCarthy and McDonald 2003), but its scalability and computational potency is limited by the Windows platform. LTR_finder (Xu and Wang 2007) and LTRharvest (Ellinghaus, et al. 2008) are by far the most sensitive programs in finding LTRs. Nevertheless, these programs suffer from reporting large numbers of false positives (Lerat 2010). MGEScan-LTR is another early development of LTR searching programs (Rho, et al. 2007). Its recent update on the web-based platform allows wider usage (Lee, et al. 2016), but is still associated with the issue of false identifications. As the most sizeable content of plant genomes, the assembly of LTR-RTs in plant genomes is typically compromised due to the collapse of short reads from such regions. Fragmented and misassembled repetitive sequences could lead to further error propagation in downstream genome annotation. Unfortunately, most of the current programs are not well adapted to the nature of draft genomes.

In this study, we introduce LTR_retriever, a novel tool for identification of LTR-RTs. This package efficiently removes false positives from initial software predictions. We benchmarked the performance of LTR_retriever with existing programs using the well assembled and annotated rice genome (International Rice Genome Sequencing Project 2005) as well as other high-quality monocot and dicot model genomes, e.g., maize (Jiao, et al. 2017), sacred lotus (*Nelumbo nucifera*) (Ming, et al. 2013), and Arabidopsis (*Arabidopsis thaliana*) (Arabidopsis Genome Initiative 2000). Our results indicated that LTR_retriever achieved very high specificity, accuracy, and precision without significantly sacrificing sensitivity, hence significantly outperforming existing methods. In addition, we implemented a module to accurately search for non-canonical LTR-RTs that featured non-TGCA motifs in LTR regions. A search in 50 published genomes identified seven types of non-canonical LTR-RTs, which are mainly *Copia* elements with substantially shorter length compared to regular *Copia* elements. Further characterizations show that non-canonical LTR-RTs are less abundant in the genomes but preferentially inserted into genic regions. Finally, we demonstrated the feasibility of making high-quality LTR libraries from self-corrected PacBio reads.

## NEW APPROACHES

*De novo* prediction of LTR-RTs can produce large amounts of false positives. To detect and filter out non-LTR sequences and obtain high-quality LTR-RT exemplars (representative LTR-RT sequences), we developed eight modules with adjustable parameters in LTR_retriever (**Fig 2**). A detailed description of each individual module can be found in **Supplementary Methods**.

**Fig 2.**
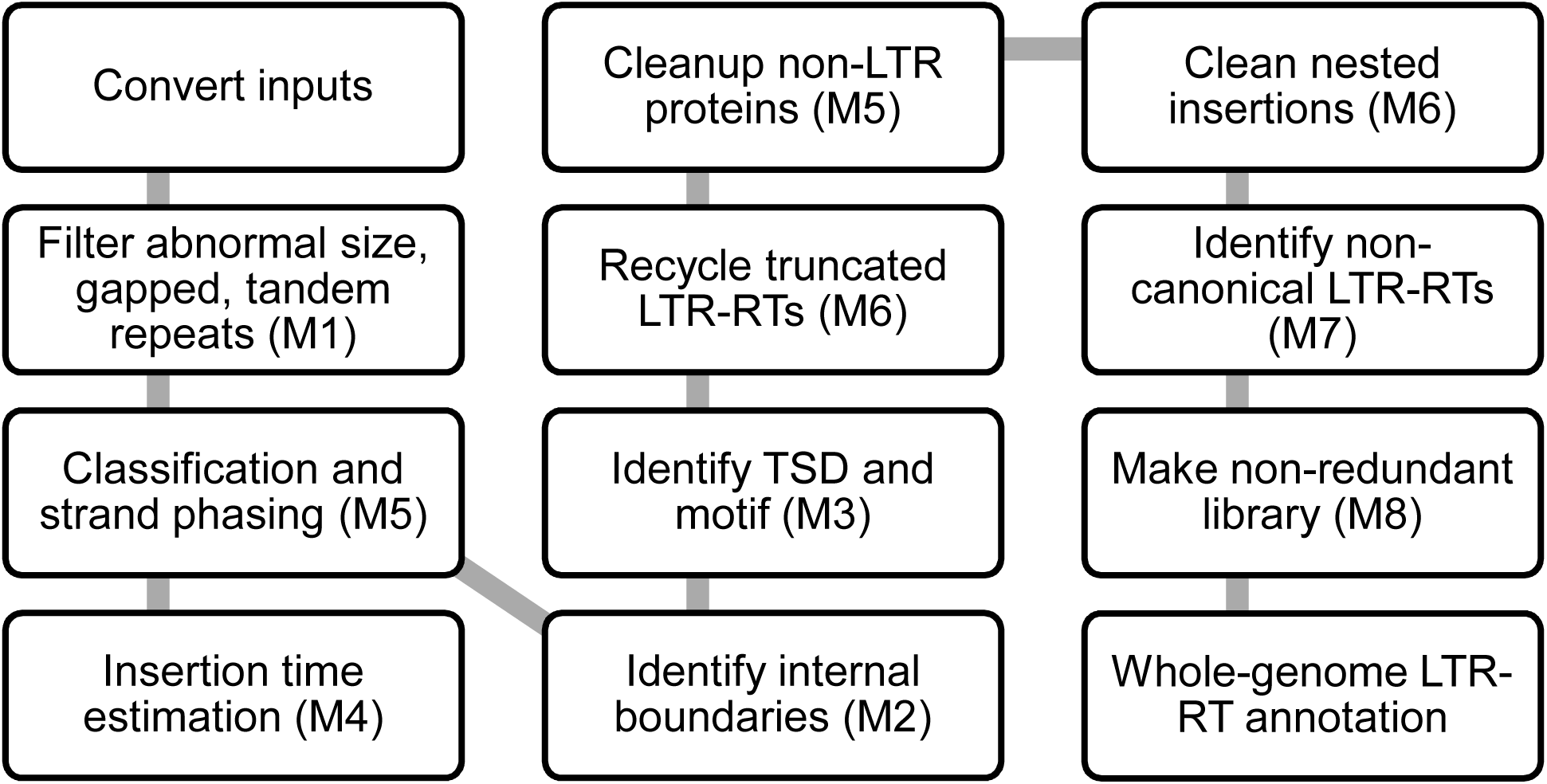
Workflow of LTR_retriever. Modules 1-8 are indicated in parentheses.

## RESULTS

Recovery of LTR elements based on structural features has been implemented in multiple packages. However, high level of false positive is a key issue. It is possible to reduce false positives by defining more stringent parameters such as high LTR similarity, intermediate LTR length, and “TGCA” motif (**Fig 3, Supplementary Table S1**). Unfortunately, the level of false negatives becomes high when more stringent parameters are applied (**Fig 3, Supplementary Table S1**). The trade-off between sensitivity and specificity cannot be minimized by merely adjusting parameters of existing tools (**Fig 3, Supplementary Table S1**). To establish efficient filters, it is essential to understand the fundamental differences between true LTR elements and false positives. In this study, we employed four statistical metrics (sensitivity, specificity, accuracy, and precision) to evaluate the performance of LTR-RT recovery programs (**Materials and Methods**).

**Fig 3.**
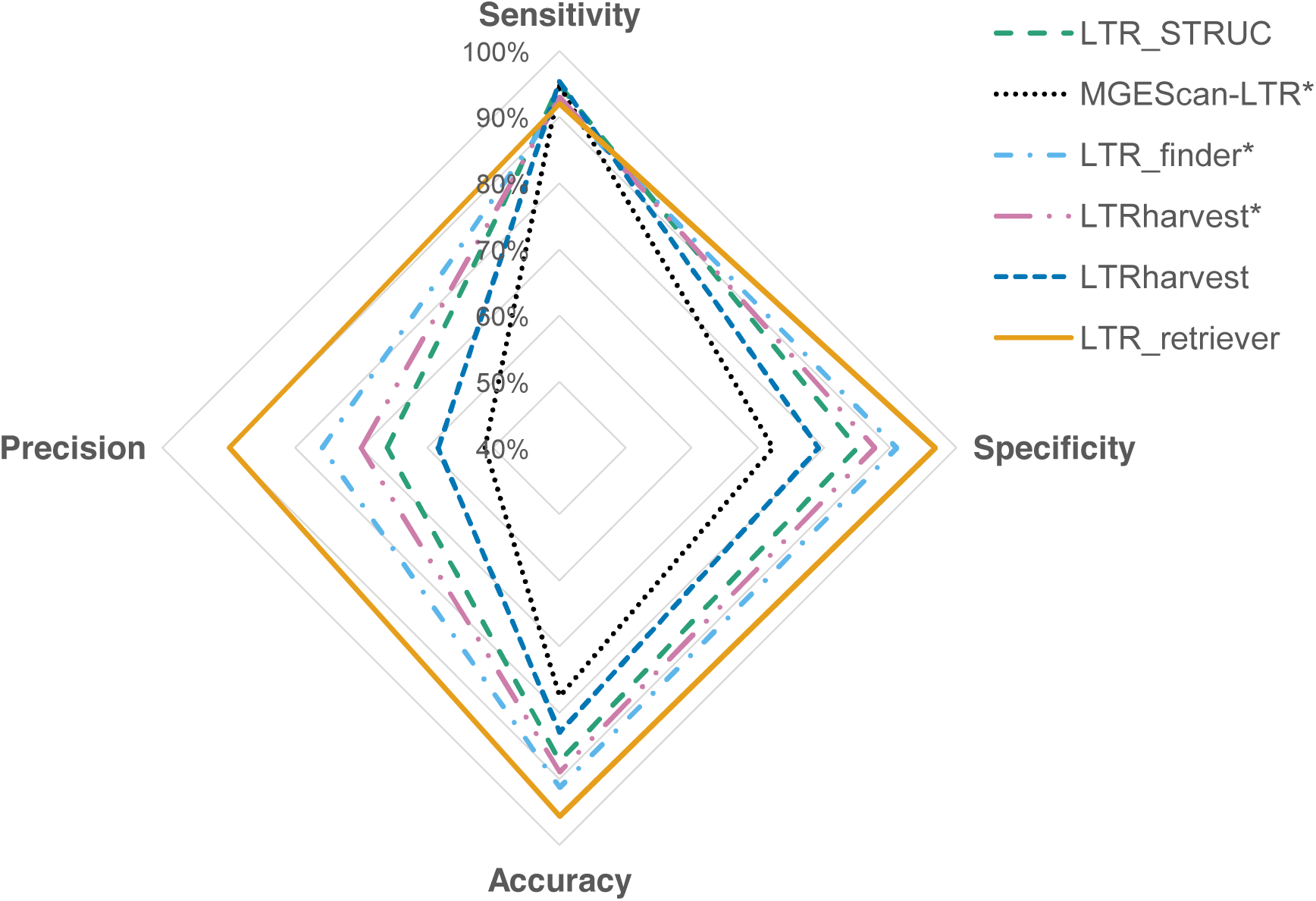
Comparison of the performance of LTR-RT recovery programs on the rice genome. LTR libraries of the rice genome were constructed using LTR_STRUC, MGEScan-LTR, LTR_finder, LTRharvest, and LTR_retriever, respectively, and then were used to identify LTR sequences in the genome using RepeatMasker. Identified candidate sequences were compared to whole-genome LTR sequences recognized by the manually curated standard library. The genomic size (bp) of true positive, false positive, true negative, and false negative were used to calculate sensitivity, specificity, accuracy, and precision. *Indicates the analysis were using optimized parameters (**Materials and Methods**) while the remainder was in default parameters.

### Features of LTR false positives and solutions

In genome assembling practices, one of the most difficult tasks is to assemble highly repetitive regions. Even in the best-assembled genomes, there are still gaps to be filled. In assemblies of non-overlapping scaffolds, sequence space (gaps) is manually added based on their inferred order. For a piece of sequence with gaps, it is not uncommon that genome assemblers mistakenly join two similar sequences that belong to different transposable elements from the same family. Under these situations, the ambiguous sequence replaced by gaps is much less reliable than continuous sequence.

Tandem repeats are locally duplicated sequences of two or more bases such as centromere repeats and satellite sequences (Benson 1999). Although it is possible that an LTR element carries small portions of tandem repeats, it becomes an LTR false positive when the majority sequence of an LTR-RT candidate consists of tandem repeats including low complexity sequences. We deploy **Module 1** in LTR_retriever to eliminate candidates that contain substantial amounts of gaps and tandem repeats (**Fig 2, Supplementary Methods**). **Module 1** also controls sequence length in consideration of both extremely long (15KB) and short (100bp) LTR-RT. The broad range of length settings allows LTR_retriever to identify very short elements like TRIM or exceptionally long elements. The implementation of **Module 1** allows LTR_retriever to exclude 4~12% of total candidates which are very likely false positives.

Identifying the exact boundaries of an LTR candidate is critical for further structural analysis such as motifs and TSDs. Published methods have applied some schemes to define boundaries. In practice, we found that the external boundaries of an LTR candidate were defined quite precisely by these prediction methods. However, for the internal boundaries which define the start and end of the internal region, predictions of existing methods are often incorrect. By manual inspections, we found the percentage of inaccurate internal boundary could be as high as 30%. The misdefined internal boundary of an LTR candidate will result in an incorrect prediction of LTR structures, such as motif, PBS, and PPT, which is likely to fail in the next filtering steps. We thus developed **Module 2** for correction of the internal boundaries of raw LTR predictions (**Fig 2, Supplementary Methods**), which could recover an extra 27% high-quality LTR candidates in the rice genome.

LTR-RT features with long terminal repeat flanking each side of the internal region. To exhaustively search for LTR candidates from genomic sequences, most published tools start with finding sequence alignments that are close to each other. This approach can effectively identify LTR elements featured with a pair of long terminal repeats as well as finding non-LTR TE pairs that are similar to each other (**Fig 1**). Such non-LTR TE fragments could be contributed by tandem repeats, DNA TEs, SINEs, LINEs, solo-LTRs from the same LTR-RT family, or other repetitive sequences including tandemly located gene families. Excluding such LTR-like false positives is challenging.

Moreover, consider that some TEs prefer to insert into other TE sequences, TE clusters are frequently found (SanMiguel, et al. 1998; Bergman, et al. 2006). The dense distribution of TEs creates a significant amount of false LTRs in *de novo* predictions. With close inspection, we found that in most cases, the intra-element sequence similarity of such false positives extended beyond the predicted boundary of the direct repeat (**Fig 1E**). In contrast, for a true LTR-RT, the sequence alignment terminates at the boundary of the LTR region. This represents an important structural feature that could distinguish LTR-RTs and its false positives. Another distinctive feature between true LTR and such false positives is the existence of TSDs. In an LTR-RT, TSDs flanking the element are identical (**Fig 1A**). However, in an LTR false positive, sequences at each end have different origins (**Fig 1E**). For 4-6 bp random sequences, the possibility of one being identical to the other is 0.02-0.39%, which is very unlikely. To utilize the structural difference between LTR-RT and false positives, **Module 3** was developed (**Fig 2, Supplementary Methods**) to exclude elements with extended alignment beyond LTR regions and those without a TSD immediately flanking the termini of LTRs. Benefiting from the accurate boundaries of candidate elements corrected by **Module 2**, this module could effectively identify most of the false positives which could account for nearly half (42.6%) of total LTR candidates.

**Module 3** also allows fine-grained adjustment of the internal and external element boundaries by jointly searching TSDs and motifs. As LTR-RTs are predominantly represented by 5 bp TSD and the 5'-TG..CA-3' motif, searching for such sequence structure at the termini of direct repeats is prioritized. If the canonical motif is absent, the seven non-canonical motifs (**Supplementary Table S2**) is searched instead. This function allows LTR_retriever flexibly while accurately characterizing the terminal structure of an LTR candidate. In rice, up to 99% of recognized LTR-RTs carry the canonical 5'-TG..CA-3' motif immediately flanked by 5 bp TSDs, while less than 0.1% of LTR-RTs have non-canonical motifs with 5 bp TSDs. In other cases, LTR candidates were found carrying the canonical motif with TSDs less than 5bp, which could be due to inter-element recombination or mutation. For example, in the maize genome, LTR-RT with TSD length of 3 bp and 4 bp have 108 and 483 occurrences out of 43,226 intact LTR-RTs, respectively.

Similar to retroviruses, direct repeats of a newly inserted LTR-RT are identical to each other. Based on the neutral theory (vonHoldt, et al. 2012), **Module 4** was developed for the estimation of insertion time of each intact LTR-RT (**Fig 2, Supplementary Methods**). We applied the Jukes-Cantor model for estimation of divergence time in noncoding sequences (Jukes and Cantor 1969). In the rice genome, more than 99% of intact LTR-RTs are inserted within 4 million years (MY) given the rice mutation rate of 1.3 × 10^-8^ mutations per site per year (Ma and Bennetzen 2004) (**Supplementary Fig S2**).

In the internal region of an LTR element, coding sequences like *gag*, *pol*, and *env* are usually found (**Fig 1A**) (Ellinghaus, et al. 2008), which could also help to discriminate LTR-RTs and non-LTRs efficiently. In **Module 5**, we applied the profile hidden Markov model (pHMM) to identify conserved protein domains that occur in LTR-RT candidate sequences (**Fig 2, Supplementary Methods**). A total of 102 TE-related pHMMs were identified using the rice TE library, with 55 non-LTR profiles and 47 LTR-RT profiles which include 30 *Gypsy* profiles, 9 *Copia* profiles and 8 profiles with ambiguous LTR-RT family classifications (unknown). In rice, 82.6% of intact LTR-RTs could be classified as either *Copia* or *Gypsy* using **Module 5**. Furthermore, the direction of LTR-RT could be phased using the profile match information. Eventually, 60.5% of LTR-RTs in rice could be phased to either on the positive strand or negative strand. A BLAST-based search for non-LTR transposase and plant coding proteins in LTR-RT candidates are also implemented in **Module 5** for the further exclusion of non-LTR contaminations. About 1-4% of the candidate sequences were recognized as non-LTR originated and could be further eliminated.

After screening and adjustment of LTR candidates using **Module 1** to **Module 5**, the retained candidates are structurally intact LTR-RTs. However, since the screening criteria are very stringent, some true LTR-RTs could be excluded. Through manual inspection, we found that some LTR-RT candidates passed all the screening criteria but only have minor deletions at either the 5' or 3' termini, resulting in the failure in the identification of terminal structures. Such candidates are categorized as truncated LTR-RTs whose intact LTR region and the internal region will be retained if there is no highly similar copy in the intact LTR element pool. **Module 6** was designed to retain sequence information from truncated LTR-RTs which contributes about 10% of sensitivity increment of LTR_retriever (**Fig 2, Supplementary Methods**).

New LTR-RT tends to insert into other LTR-RTs, creating nested insertions. To exclude nested insertions from the LTR exemplars, we developed a function in **Module 6**, which utilizes all newly identified LTR regions to search for homologous sequences in identified internal regions. This search could recognize and removes LTR-RTs that are nested in intact LTR-RTs. Using this method, about 8% of LTR-RT internal regions in rice and 67.7% in maize are identified as nested with other LTR elements. By removing such nested insertions, the library size can be reduced significantly without sacrifice of sensitivity. More importantly, it avoids the misannotation of LTR sequences as internal regions.

### Construction of non-redundant LTR library

Construction of the repeat library with non-redundant, high-quality TE sequences is critical for RepeatMasker-based TE and gene annotations, with the size of the repeat library being one of the limiting factors for speed. The required time for whole genome TE annotations using RepeatMasker is highly correlated to the size of TE libraries. Since the identified LTR-RTs are redundant, it would significantly speed up whole genome LTR-RT annotation if the redundancy is eliminated. To reduce redundancy of identified LTR-RTs, **Module 8** was developed using the clustering function of BLAST or CD-HIT. Due to the reduced redundancy and exclusion of nested insertions (**Module 6**), the LTR-RT sequence size was reduced to 10-30% of its original size. Accordingly, whole genome LTR-RT annotation could be accelerated ∼4-fold with similar sensitivity comparing to a non-redundant LTR library.

### Comparison of performances to other LTR identification tools

To compare the performance between LTR_retriever and other existing methods, we employed the rice genome as a reference. The rice genome is one of the best sequenced and assembled genomes (International Rice Genome Sequencing Project 2005). To set a standard for our comparison study, we manually curated representative LTR elements obtained from the rice genome (cv. Nipponbare) and generated a compact repeat library which contains 897 sequences with the size of 2.34 Mb. The 897 sequences represent 508 non-redundant LTR elements (**Supplementary Methods** and **Supplementary Sequence Files**). Using this library, LTR-RT contributes 23.5% of the assembled genome (374 Mb). This number is slightly higher than the two highest estimates from previous studies (20.6%, 22%) (Ma, et al. 2004; Chaparro, et al. 2007), suggesting the current identification of LTR retrotransposon in Nipponbare is close to saturation and the library is reasonably comprehensive. As a result, this library is used as a reference library for subsequent analysis. The accurate annotation of LTRs in the rice genome allows us to summarize the true positive (TP), true negative (TN), false positive (FP), and false negative (FN) of a *de novo* LTR prediction and annotation, hence allowing the evaluation of different methods.

The sensitivity of all existing LTR discovery tools was reported very high (Xu and Wang 2007; Ellinghaus, et al. 2008; You, et al. 2015), however, systematic evaluation of specificity using the whole genome sequence length is not available. Specificity describes the proportion of true negative, i.e., non-LTR sequences, being correctly ruled out, which is as important as sensitivity for evaluation of a diagnostic test (Zhu, et al. 2010). To better describe the performance of these methods, precision and accuracy are also calculated (Fawcett 2006). Precision, or positive predictive value, is the proportion of true positives, i.e., LTR sequences, among all positive results revealed by the test. The precision is an indication of false discovery rate (FDR), with the equation FDR=1-precision. Accuracy is the proportion of true predictions, which controls systemic errors and random errors (**Materials and Methods**).

For comparison, we chose four of the most widely used LTR searching methods, LTR_STRUC (McCarthy and McDonald 2003), MGEScan-LTR (Rho, et al. 2007), LTR_finder (Xu and Wang 2007), and LTRharvest (Ellinghaus, et al. 2008), for performance benchmarks. As LTRharvest is the most flexible program with more than 20 modifiable parameters, we optimized the parameters based on our experience for more accurate predictions (**Fig 3**). The optimized parameters were also applied to the parameter settings of LTR_finder and MGEScan-LTR. LTR_retriever can utilize multiple input sources including the results from LTR_finder, LTRharvest, and MGEScan-LTR. We used separate and combined inputs in LTR_retriever for comparisons.

As expected, sensitivities of the most published methods are very high, ranging from 91.2% to 95.3% (**Fig 3, Supplementary Table S1**). However, specificities of these methods are not desirable, ranging from 72.3% to 87.7% (**Fig 3, Supplementary Table S1**) with the exception of LTR-finder (91.0%). Specificity of 72.3% indicates that 27.7% of non-LTR genomic sequences were falsely recognized as LTR-RT sequences. The optimized parameters in LTRharvest led to an improvement of the specificity from 79.2% to 87.7% (**Supplementary Table S1**). The optimized LTR_finder had the best balance, with sensitivity and specificity both reached to the level of 90%, however, its precision is only 75.8% (**Fig 3, Supplementary Table S1**). As a reminder, FDR=1-precision. Although LTR_finder has the highest precision among the published methods, the precision of 75.8% indicates that 24.2% of “LTR-RT related sequences” identified in the genome were falsely reported as LTR-RT. The accuracy of existing methods ranges from 77.5-91.3%, showing variations in true prediction rate.

We tested LTR_retriever using the optimized LTRharvest results as input. As a stringent filter, LTR_retriever achieved specificity and accuracy of 96.8% and 95.5%, respectively, greatly outperforming existing methods (**Fig 3, Supplementary Table S1**). The precision also increased from the original 69.9% to 89.9%, indicating the FDR dropped to 1/3 and is among the lowest of all methods (**Fig 3, Supplementary Table S1**). Strikingly, the sensitivity of LTR_retriever remained as high as 91.1% compared to the original 93.0%, meaning that we only sacrificed less than 2% of sensitivity to achieve the observed performance improvements (**Fig 3, Supplementary Table S1**). Other input sources such as those from LTR_finder and MGEScan-LTR were also tested and showed excellent performance (**Supplementary Table S1**). Upon combination of two or more input sources, the sensitivity is increased to 94.5%, which is equivalent to the highest level that was achieved by the existing methods, providing a workaround to achieve comprehensive and high-quality predictions (**Supplementary Table S1**). By excluding the majority of false positives, the final library size was substantially reduced, from the largest 44.4 MB by MGEScan-LTR to the final 4.4 MB by the LTR_retriever (**Supplementary Table S1**). The reduced library size significantly reduced the annotation time using RepeatMasker.

### Benchmarking on other genomes

LTR_retriever was developed based on the rice genome, which has demonstrated the highest specificity, accuracy, and precision among its counterparts with the same level of sensitivity. To test whether the excellent performance of LTR_retriever can be reproduced with other genomes, we chose four other genomes with variable amounts of LTR elements including two maize genomes (cv. B73 and cv. Mo17) (Xin, et al. 2013; Jiao, et al. 2017), Arabidopsis (Arabidopsis Genome Initiative 2000), and sacred lotus (Ming, et al. 2013). All these genomic sequences are associated with reasonable repeat libraries so that performance of LTR_retriever could be evaluated by comparisons between the respective standard annotations and LTR_retriever generated libraries.

For all the genomes we tested, LTR_retriever demonstrated very sensitive and accurate performance in retrieving LTRs. Most metrics reached the levels of 90% (**Table 1**). For Arabidopsis, we obtained a very high specificity and accuracy, which were 98.9% and 98.4%, respectively, indicating the nearly perfect prediction by LTR_retriever. For the ancient eudicot sacred lotus, the four metrics ranged from 81.2% to 91.3%. The maize genome is known to be highly repetitive, and we used both the reference B73 (v4) and the Mo17 genomes to evaluate the performance of LTR_retriever. With LTR-RTs comprising ~75% of the 2.1 GB genome, LTR_retriever could identify 91.1% and 95.7% LTR-RTs with specificities of 90.6% and 95.7%, respectively. Due to the high LTR-RT content and the nearly perfect performance of LTR_retriever, the precisions reached 96.6% (FDR=3.4%) and 98.7% (FDR=1.3%), respectively. It is known that structure of the maize genome is very complex due to intensive nested TE insertions (SanMiguel, et al. 1996), LTR_retriever is able to overcome complex structures and recover most LTR-RTs from the genome.

**Table 1.**
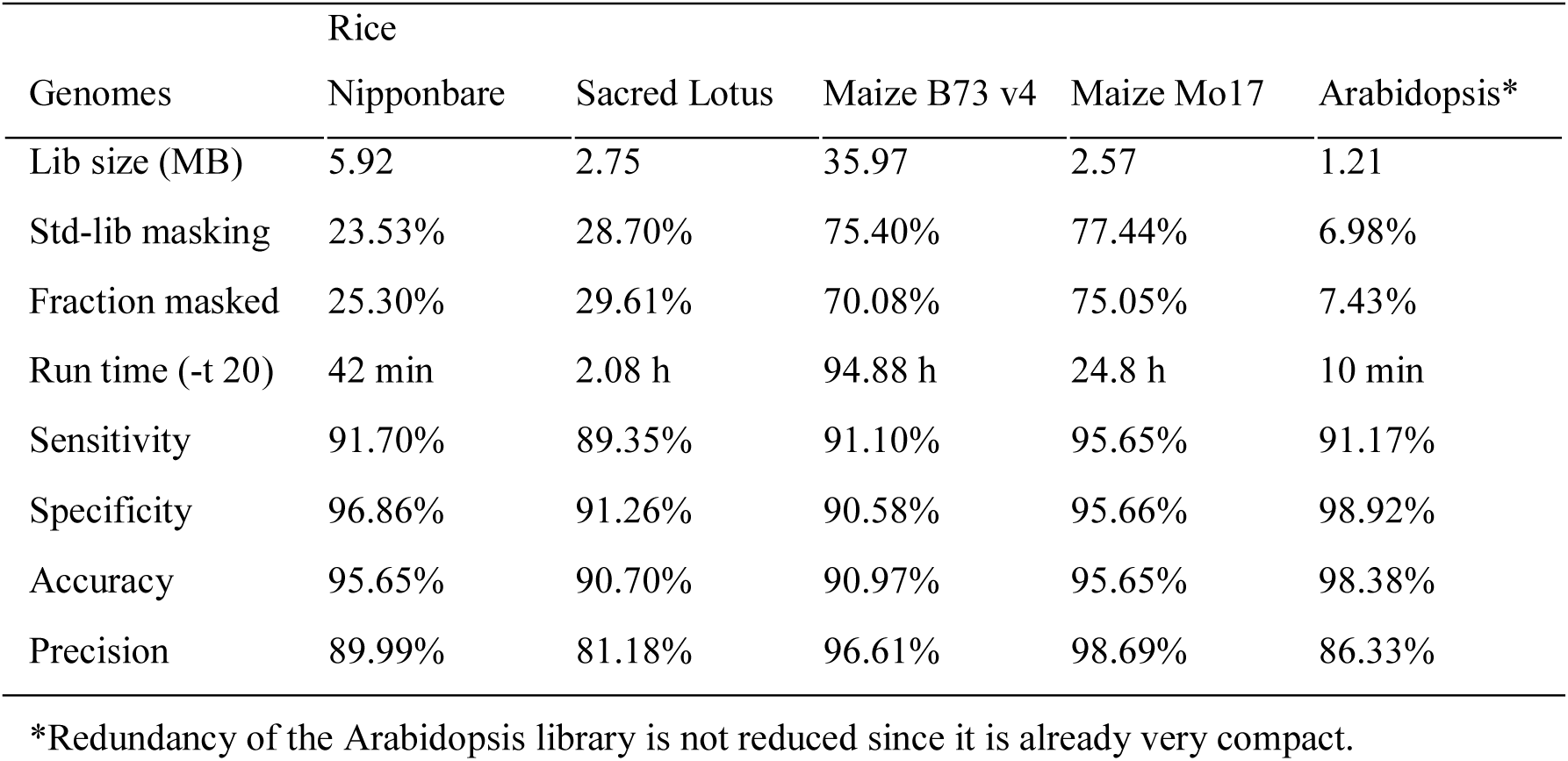
Performance of LTR_retriever on model plant genomes.

### Direct LTR library construction from PacBio reads

The recent development of long-read sequencing technologies has provided a solution for resolving highly repetitive regions in *de novo* genome sequencing projects (VanBuren, et al. 2015). The PacBio single molecule, real-time (SMRT) sequencing technology produces long reads with an average length of 10-15kb. Empirically, more than 95% of LTR-RTs range from 1-15kb (**Supplementary Fig S1**). Thus, theoretically, the long-read sequencing technology may allow us to identify intact LTR elements directly from the reads.

It is known that the current PacBio RS II platform has an average sequencing error rate of 15%. In our experience, most LTR-RT insertions are structurally detectable if inserted 4 million years ago or younger (**Supplementary Fig S2**) which is equivalent to 89.6% of identity between two LTR regions. When mutations/sequencing errors accumulated, the fine structure such as TSD and terminal motifs could be mutated and element would be beyond the detection limit. Thus the sequencing error rate of 15% could have artificially aged the actual LTR element to become undetectable. We tested the LTR_retriever using raw PacBio reads and no confident intact LTR element was reported. However, LTR_retriever performed excellently using self-corrected PacBio reads with an error rate of 2%.

To test the efficiency of LTR_retriever, we used 20 thousand (k) self-corrected PacBio reads from Arabidopsis L*er*-0 as an initial input (**Materials and Methods**), and with 20 k reads as an increment until 180 k. The Arabidopsis repeat library from Repbase was used to calculate sensitivity, specificity, accuracy, and precision. The LTR library constructed from the Arabidopsis L*er*-0 genome was used as the control to compare to the quality of LTR libraries constructed from PacBio reads. As more reads were used, the prediction of intact LTR-RTs increased linearly (**Fig 4A**). However, the size of LTR libraries constructed from these candidates are not increased at the same rate (**Fig 4A**), and the sensitivity exceeds the library developed from the genome sequence after 40 k reads input and is saturated at 93% after 120 k reads being used (**Fig 4B**). Since the average length of these reads is 14.6kb, and the Arabidopsis “L*er*-0” genome was assembled as ~131 MB, the sample of 40 k and 200 k reads is equivalent to 4.5- and 13.4-fold genome coverage, respectively. Moreover, despite the number of reads being used, the average specificity, accuracy, and precision were 99.5%, 98.8%, and 94.0%, respectively, indicating very high-quality LTR libraries could be constructed from PacBio reads. Furthermore, masking potentials (percentage of the genome that could be masked) of PacBio LTR libraries surpass the standard library level after using 40 k or more reads (**Supplementary Fig S3**), indicating that it is sufficient to construct a comprehensive library using as little as 4.5X PacBio self-corrected reads. To summarize, LTR_retriever shows high sensitivity, specificity, accuracy, and precision to construct LTR libraries directly from self-corrected PacBio reads prior to genome assembly.

**Fig 4.**
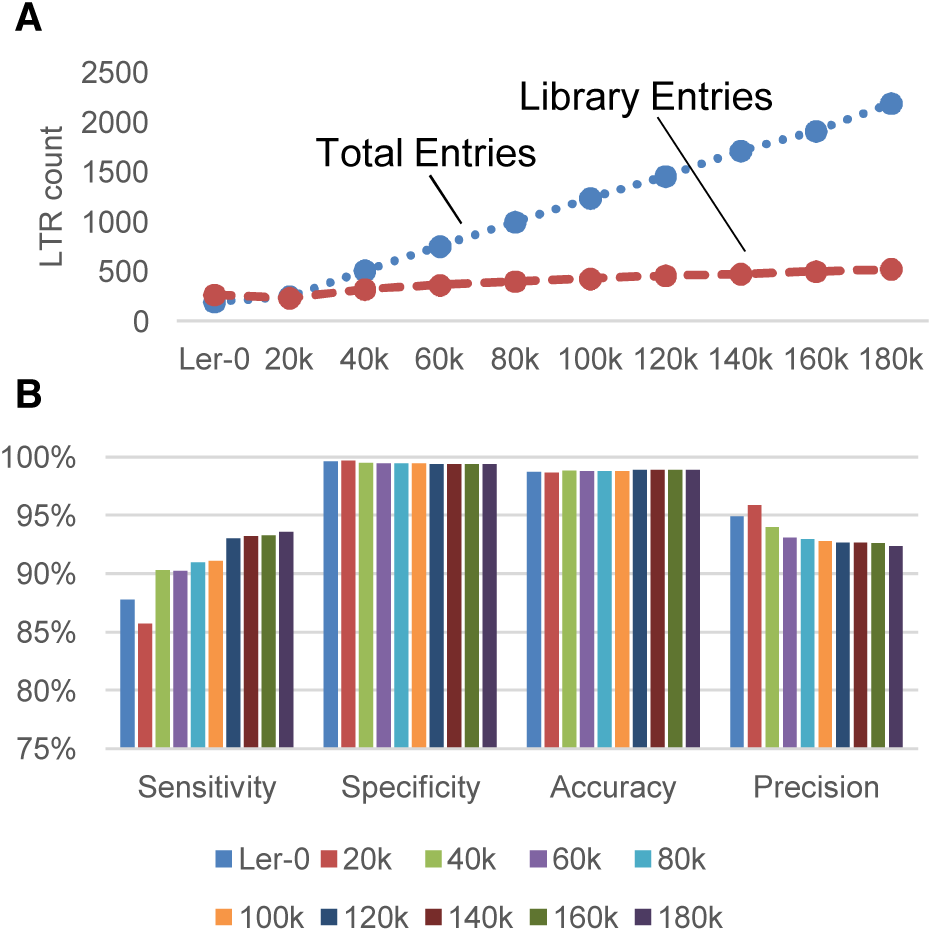
Direct library construction using self-corrected PacBio reads. (**A**) Identification of intact LTR elements and construction of libraries using the Arabidopsis “L*er*-0” genome and 20k -180k self-corrected PacBio reads. (**B**) The performance of custom LTR libraries compared with that from the Arabidopsis reference (*Col-0*) genome.

### Identification of LTR-RTs with non-canonical motifs

LTR-RT features dinucleotide motifs flanking the direct repeat regions (**Fig 1**). The most common motif is the palindromic 5'-TG..CA-3' motif. However, during manual curation of LTR-RTs, we discovered many LTRs with non-TGCA motifs (Ferguson and Jiang, unpublished). These non-canonical motifs can be non-palindromic, for example, *Tos17*, a rice LTR-RT that can be activated by tissue culture, has non-canonical motifs of 5'-TG…GA-3' (Hirochika, et al. 1996); *AtRE1* in Arabidopsis has 5'-TA…TA-3' motifs (Kuwahara, et al. 2000); and *TARE1*, intensively amplified in the tomato genome, has 5'-TA…CA-3' motifs (Yin, et al. 2013). In addition, three copies of *Gypsy*-like elements with 5'-TG..CT-3' motifs were annotated in the soybean genome (Du, et al. 2010).

To recover LTR elements with certain terminal motif, LTRharvest enables the “-motif” parameter allowing users to specify the motif to be discovered, which requires prior motif knowledge. When users apply the default setting (no motif specified), the number of LTR-RT candidates can be 2-4 times more than the result with “-motif TGCA” specified. The significant increase of predicted candidates does not necessarily indicate a large number of non-TGCA LTR recovered. With annotations and further curations, we found 99% of the additional candidates are false positives in the rice genome.

To identify non-TGCA LTR-RT with high confidence, we developed **Module 7** as an optional add-on to LTR_retriever (**Supplementary Methods**). The sacred lotus genome carries many non-canonical LTR elements. We tested the performance of LTR_retriever in identifying such elements using the manually curated non-canonical LTR-RTs from this genome (**Supplementary methods**). Our results showed that LTR_retriever could identify high-quality non-canonical LTR-RTs, with a sensitivity of 74.7% and a precision of 81.6% (FDR=18.4%). And the specificity and accuracy were 98.5% and 96.5%, respectively, indicating that the identified non-canonical LTR-RTs are highly accurate.

### Non-canonical LTR-RTs are widespread in plants and preferentially insert in genic regions

To characterize non-TGCA LTR-RTs, we searched through 50 publically available plant genomes. A total of 870 high-confidence non-TGCA LTR-RTs were found from 42 of these genomes (**Materials and methods**). Further categorization of non-TGCA LTR-RTs identified seven types of high-confident non-canonical motifs including three (TACT, TGTA, and TCCA) that were not previously reported (**Supplementary Table S2**). Further classification of ORFs within these elements based on pHMM search indicated that 89% of classified non-TGCA LTR elements were the *Copia* type, while only 11% were the *Gypsy* type (**Supplementary Table S2**). We also identified 83,368 canonical LTR-RTs in these genomes, with a *Gypsy* - *Copia* ratio of 2.9:1 (**Table 2**).

**Table 2.**
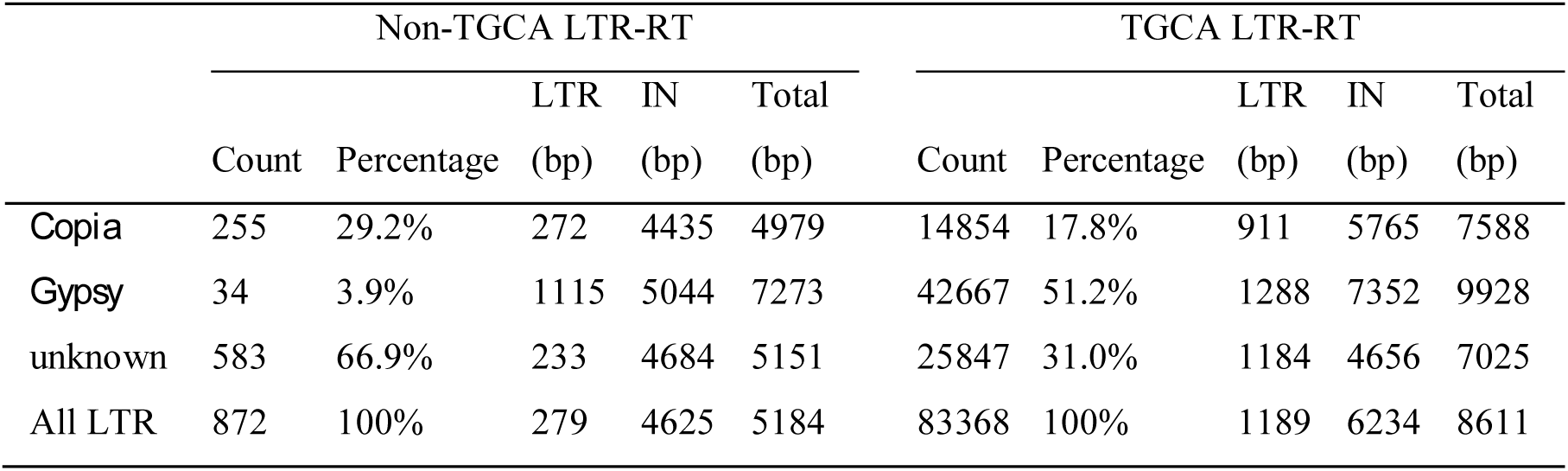
Average element size of different types of LTR-RTs in 50 sequenced plant genomes.

|  | Non-TGCA LTR-RT |  |  |  |  | TGCA LTR-RT |  |  |  |  |
| --- | --- | --- | --- | --- | --- | --- | --- | --- | --- | --- |
|  | Count | Percentage | LTR (bp) | IN (bp) | Total (bp) | Count | Percentage | LTR (bp) | IN (bp) | Total (bp) |
| *Copia* | 255 | 29.2% | 272 | 4435 | 4979 | 14854 | 17.8% | 911 | 5765 | 7588 |
| *Gypsy* | 34 | 3.9% | 1115 | 5044 | 7273 | 42667 | 51.2% | 1288 | 7352 | 9928 |
| unknown | 583 | 66.9% | 233 | 4684 | 5151 | 25847 | 31.0% | 1184 | 4656 | 7025 |
| All LTR | 872 | 100% | 279 | 4625 | 5184 | 83368 | 100% | 1189 | 6234 | 8611 |

For canonical LTR-RTs, the length of the LTR region in *Gypsy* elements is about 40% longer than *Copia* elements (**Table 2**). However, in the case of non-canonical LTR-RTs, this size difference is intensified to 400%. This is due to the significant reduction of LTR length of non-canonical *Copia* elements, from an average size of 911 bp to 272 bp (**Table 2**). The size of internal region and whole element of non-canonical *Copia* are also much shorter than those of *Copia* elements carrying the TGCA motif (**Table 2**). These results suggest that shorter LTRs may have facilitated the amplification and survival of non-TGCA LTR-RTs.

Comparing to canonical *Copia* elements, less new insertions (5% less for elements younger than 0.2 MY) and more old elements (7% more of 1.2 MY – 1.8 MY elements) (**Fig 5A**) were observed for non-canonical *Copia* elements based on sequence similarity between LTR sequences. Meanwhile, we found that elements with canonical motifs were more likely to form solo LTRs. Comparing to 54% of the non-canonical *Copia* elements have solo-complete LTR ratios less than three, only 32% of canonical *Copia* elements are in this category, indicating the inefficient removal of non-canonical LTR-RT insertions (**Fig 5B**). To characterize the insertion preference, we extracted 200 bp flanking sequences of each element, and BLAST against the genome for determination of copy numbers. The majority (70%) of the flanking sequences of non-canonical *Copia* elements have copy numbers less than five, while that of canonical *Copia* elements is 46% (**Fig 5C**). Strikingly, 40% of non-TGCA *Copia* elements are located within 1KB distance to protein-coding genes, which is 16% more frequent than canonical *Copia* elements (**Fig 5D**). Taking together, our results show that non-canonical *Copia* elements prefer non-repetitive genomic regions and are often inserted within or close to genes.

**Fig 5.**
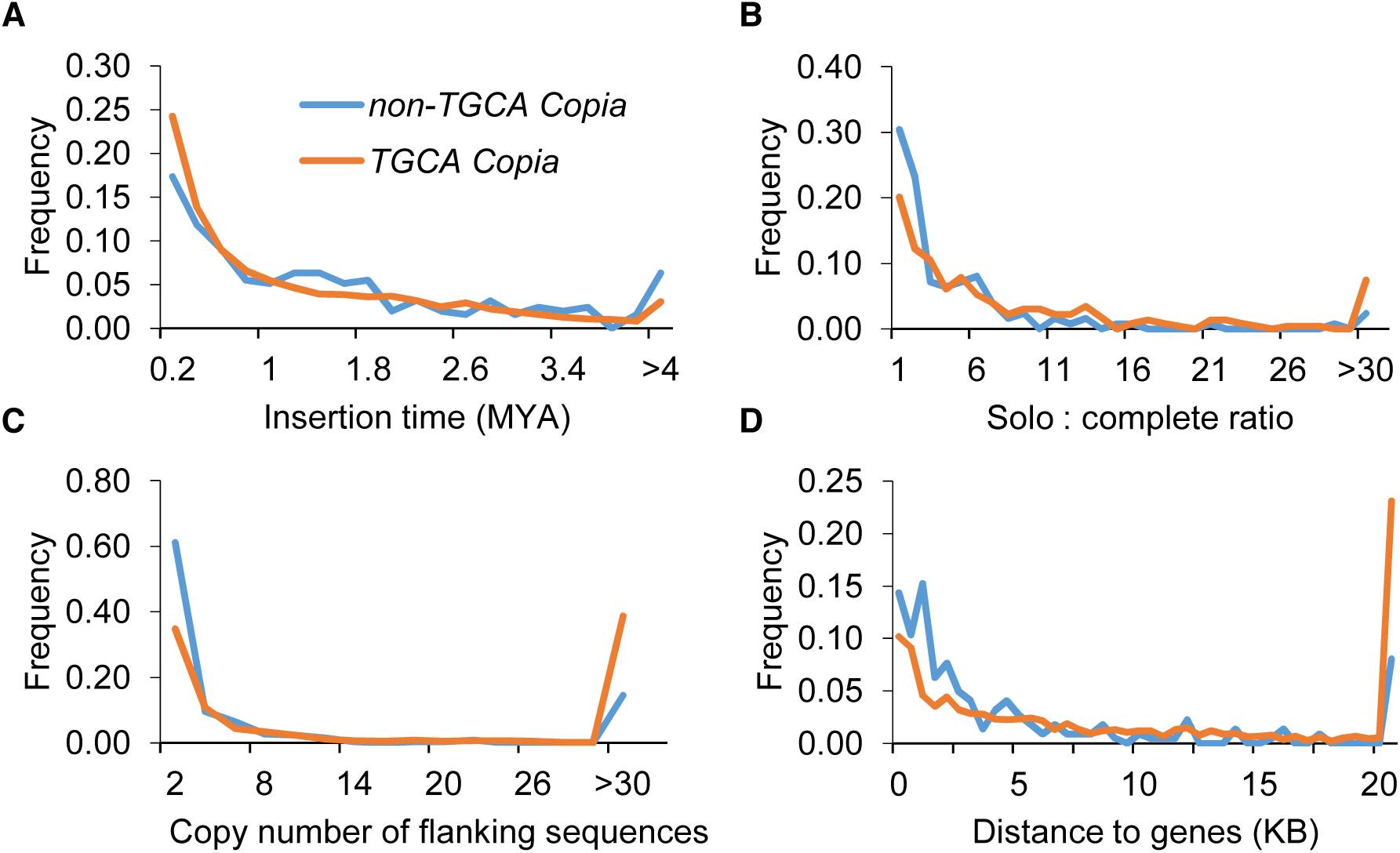
Characterization of non-canonical *Copia* elements in plants. (**A**) Non-TGCA *Copia* is older than canonical *Copia*. (**B**) Non-TGCA *Copia* has lower ratio of solo LTR to complete LTR, indicating ineffective exclusion for this type of LTR elements. (**C**) Non-TGCA *Copia* elements are predominately associated with non-repetitive flanking sequences. (**D**) Non-TGCA *Copia* elements are located closer to genes than canonical *Copia* elements. Blue lines represent non-TGCA (non-canonical) *Copia* elements and orange lines represent TGCA (canonical) *Copia* elements. All analyses were based on 50 plant genomes.

## DISCUSSION

Technological advances have minimized the cost of sequencing a genome. The real bottleneck to establishing genomic resources of an organism is the annotation of its genomic sequence. As mentioned above, TEs, particularly LTR retrotransposons, are the largest component of most plant genomes. If TEs are left unmasked prior to gene annotation, they would seed numerous of spurious sequence alignments, producing false evidence for gene identification. Even worse, the open reading frames of TEs look like *bonafide* genes to most gene-prediction software, corrupting the final annotations. As a result, the first step of genome annotation is to identify TEs and other repeats. Subsequently, these repeats are masked to facilitate gene annotation. As a result, the quality of repeat library is not only important for the study of repeats, but also critical for high-quality gene prediction.

In this study, we reported the development of LTR_retriever, a multithreading empowered Perl program that can process LTR-RT candidates from LTR_finder, LTRharvest, and MGEScan-LTR and generate high-quality and compact LTR libraries for genome annotations or study of transposable elements. We curated LTR elements identified from the rice genome and used the curated LTR library as the standard to test the performance of LTR_retriever in terms of sensitivity, specificity, accuracy, and precision. Benchmark tests on existing programs indicated very high sensitivities achieved, however, specificities and accuracies were not satisfactory, and the FDR could be as high as 49%, suggesting the necessity for improvement (**Supplementary Table S1**).

Since annotation of TE sequences usually precedes the annotation of functional genes for a newly sequenced genome, propagation of false positives in the construction of LTR library will significantly increase the probability of misidentification of LTR sequences in the genome and further dampen the power of downstream annotations. For example, it is known that most DNA transposons target genic regions and avoid repetitive sequences (Feschotte and Pritham 2007; Han, et al. 2013). As a result, it is not uncommon that the sequence between two adjacent DNA transposons represents gene coding regions or regulatory sequences. If the two DNA transposons are mistakenly annotated as the LTR of an individual LTR-RT, the intervening genes would be considered as the internal region of an LTR-RT and would be masked before gene annotation. In this scenario, the false positives could be extremely detrimental for downstream analyses. LTR_retriever effectively eliminates such false positives. By processing LTR-RT candidates using LTR_retriever, the specificity and accuracy reached to 96.9% and 95.7%, respectively, and the FDR is reduced to 10% which is among the lowest of all existing methods (**Fig 3, Supplementary Table S1**). Strikingly, the sensitivity of LTR_retriever remained as high as 91.7%, meaning that we only sacrificed less than 2% of sensitivity to achieve all these performance improvements (**Fig 3, Supplementary Table S1**). Further benchmark tests on two maize genomes, the sacred lotus genome, and the Arabidopsis genome also showed excellent performance (**Table 1**), suggesting that LTR_retriever is compatible with both monocot and dicot genomes.

The majority of LTR-RTs we identified carried a palindromic dinucleotide motif flanking each direct repeat. The motif is well conserved and is usually 5'-TG..CA-3'. However, the importance of such conservation is poorly understood. Retrovirus, e.g., HIV-1, is thought to be the close relative of LTR elements with the addition of an envelope protein (Zhou, et al. 2001; Hobaika, et al. 2009). Studies of retrovirus integration indicated that the terminal sequences of retroviral LTR regions, especially the 3' CA ends, are essential and important for integration of the virus (Zhou, et al. 2001; Hobaika, et al. 2009). As a result, there might be a convergent evolution between the termini of the elements and transposition machinery. That may explain why most LTR elements have the conserved TG..CA motif.

Despite the conservation, non-TGCA motifs were also found but in a much lower frequency. LTR_retriever also demonstrated high performance in identifying such non-canonical LTR-RTs. A broad scan on 50 published plant genomes retrieved seven non-TGCA type LTR-RTs with the majority belonging to the *Copia* family (**Supplementary Table S2**). For some, the abundance is not ignorable. It appears that, among the four terminal nucleotides (TGCA), only the first nucleotide (T) is invariable. Our systemic survey for the presence of non-canonical termini provides guidance for future annotation of LTR elements.

Previous studies indicate that *Gypsy* and *Copia* elements are differentially located in plant genomes. The distribution of *Copia* elements is biased toward euchromatic chromosomal arms that are relatively close to genes, whereas *Gypsy* elements are more likely located in the gene poor, heterochromatic or pericentromeric regions (Baucom, et al. 2009; Bousios, et al. 2012). Here we demonstrate, the non-canonical *Copia* elements are even closer to genes than canonical *Copia* elements and preferentially insert into non-repetitive sequences (**Fig. 4**). Apparently, there is a negative correlation between distance to genes and elements size, particularly the size of LTRs. As a result, the limited amplification and smaller size are likely the consequences of the target specificity of non-canonical LTR elements.

In Arabidopsis, TEs are separated into two classes based on their location (Sigman and Slotkin 2016). One class is present in large constitutive heterochromatic regions and their CHH methylation is maintained by chromomethylase 2 (CMT2), and the other class is located near genes where CHH methylation is constantly targeted by RNA-directed DNA methylation (RdDM). TEs in genic regions are subject to more stringent epigenetic control and demonstrate a higher level of CHH methylation compared to TEs in the non-genic region (Gent, et al. 2013; Li, et al. 2015). Moreover, TE insertions in genic regions are less likely to spread in the population since some of them are deleterious. In addition, genic space in a genome is limited comparing to the non-genic sequence space. The combined effect of epigenetic control, negative selection, and limited target sites is attributed to the low abundance of non-canonical LTR elements. Furthermore, selection against insertion of large size TEs would result in the relatively small size of both LTR and internal region of these elements. To this notion, the *Tos17* element in rice (with a “TG..GA” terminal motif) is an excellent example. The length of the *Tos17* element is only 4.3 kb with an LTR of 138 bp, which is very small compared to other autonomous LTR elements (**Table 2**). It preferentially inserts into genic regions and may amplify rapidly during tissue culture (Miyao, et al. 2003). Nevertheless, there are only a few copies of *Tos17* in the natural population of rice (Hirochika, et al. 1996), suggesting the selective pressure against insertion of this element (Hirochika, et al. 1996; Miyao, et al. 2003). Because of its insertion preference, *Tos17* has been applied as a tool for mutagenesis (Miyao, et al. 2003). In our study, we identified 870 high-confidence non-canonical LTRs in 42 out of 50 plant genomes, which is likely an underestimate due to high stringency. These elements also prefer genic insertions, which could contain other *Tos17-like* active elements in these species. In conclusion, annotation of non-canonical LTR elements is important not only due to their prevalent distribution, but also the potential application in functional studies in plants.

The recent development of single molecule sequencing technology enables the assembly of low complexity and repetitive regions. Many genome sequencing projects have benefited from the PacBio SMRT sequencing technique which features with 10-15kb average read length (Ming, et al. 2015; VanBuren, et al. 2015). Given the length of most LTR elements is less than 15kb (**Supplementary Fig S1**), it is possible to identify full-length LTRs from PacBio long reads. We applied LTR_retriever on self-corrected PacBio reads which proved a successful strategy to identify LTR-RTs. For the Arabidopsis “L*er*-0” genome, 40 thousand self-corrected reads covering approximately 4.5X of the genome were more than sufficient to generate an LTR library with higher quality compared to that generated from the assembled genome (**Fig 4**). Although self-corrected reads still have ~2% sequencing error rate, the generated LTR library was proven highly sensitive and accurate (**Fig 4**).

The pre-identified full-length LTRs may help to estimate LTR percentages of the new genome, study the evolution of LTR-RTs without performing the computationally intensive whole genome assembly, and facilitate downstream *de novo* gene annotation. Since LTR-RTs contribute greatly to the size of plant genomes, identification and masking of repetitive sequences in advance could speed up the genome assembly by as much as 50-fold (Gregory Concepcion, Pacific Bioscience, personal communication).

In summary, we developed a package which takes genome sequences or corrected PacBio reads as input and generates high-quality, non-redundant libraries for LTR elements. It also provides information about the insertion time and location of intact LTR elements in the genome. This tool demonstrates significant improvements in specificity, accuracy, and precision while maintaining the high sensitivity compared to existing methods. As a result, it will facilitate future genome assembly and annotation as well as enable rapid comparative studies of LTR-RT dynamics in multiple genomes.

## MATERIALS AND METHODS

### Implementation of LTR_retriever

LTR_retriever is a command line program developed based on Perl. The package supports multi-threading, which was achieved using the Semaphore module in Perl, and multithreading requests are passed to dependent packages. LTR_retriever takes genomic sequences in the FASTA format as input. The program can handle fragmentized and gapped regions, which is a benefit when annotating draft genomes. LTR_retriever has been optimized for plant genomes; however, its parameters can be adjusted for the genomes of other organisms. The output of the program contains a set of high-quality, comprehensive but non-redundant LTR exemplars (library), which can be used to identify or mask LTR sequences using RepeatMasker. Additionally, a summary table that includes LTR-RT coordinates, length, TSDs, motifs, insertion time, and LTR families is produced. The program also provides gff3 format output, which is convenient for downstream analysis.

### Genomes and sequences

The initial BAC sequences of “Nipponbare” were downloaded from the Rice Genome Research Program (http://rgp.dna.affrc.go.jp) for our early efforts to construct the rice TE library. The rice reference genome “Nipponbare” release 7 was downloaded from the MSU Rice Genome Annotation Project (http://rice.plantbiology.msu.edu) (Kawahara, et al. 2013). The sacred lotus genome was downloaded from the National Center for Biotechnology Information (NCBI) under the project ID “AQOG01”. The Arabidopsis reference genome “Columbia” version 10 was downloaded from The Arabidopsis Information Resource (TAIR) (www.arabidopsis.org) (Berardini, et al. 2015). The maize genome “B73” version AGPv4 was downloaded from Ensembl Plants release 34. An additional of 46 plant genomes were downloaded from Phytozome v11 (Goodstein, et al. 2012) (**Supplementary Methods**).

The Arabidopsis “L*er*-0” genome was sequenced and assembled by Pacific Biosciences using the PacBio RS II platform and the P5-C3 chemistry. The assembly is about 131 MB with a contig N50 6.36 MB (https://github.com/PacificBiosciences/DevNet). A total of 184,318 self-corrected reads were also downloaded, which is about 2.69 GB with an average read length of 14.6kb and sequence error rate < 2%, covering 20.58 X coverage of the genome.

### Standard LTR libraries

In this study, LTR libraries from four genomes (rice, maize, Arabidopsis, and sacred lotus) were used to evaluate the performance of LTR_retriever as well as existing tools. The TE database of maize was downloaded from the Maize TE database (http://maizetedb.org). The Arabidopsis repeat library athrep.ref was downloaded from Repbase (Jurka 2000). The LTR libraries for rice and sacred lotus were manually curated in the Jiang Lab (**Supplementary Methods, Supplementary sequence files**).

### Benchmark programs and parameters

LTR_STRUC (McCarthy and McDonald 2003) was obtained from Mr. Vinay Mittal via personal communications. No parameter settings were available for LTR_STRUC. LTRharvest (Ellinghaus, et al. 2008) is part of the GenomeTools v1.5.4 (Gremme, et al. 2013). Parameters for running LTRharvest were empirically optimized with “*-minlenltr 100 -maxlenltr 7000 -mintsd 4 -maxtsd 6 -motif TGCA -motifmis 0 -similar 90 -vic 10 -seed 20*”. Optimized parameters were also applied to MGEScan-LTR (Rho, et al. 2007) and LTR_finder (Xu and Wang 2007). The modified version of MGEScan-LTR was obtained from the DAWG-PAWS package (Estill and Bennetzen 2009) and was run with parameter settings “*-min-mem=20 -mim-dist=1000 -max-dist=15000 -min-ltr=50 -max-ltr=7000 -min-orf=200*”. LTR_finder v1.0.6 was run with parameter settings “*-D 15000 -d 1000 -L 7000 -l 100 -p 20 -M 0.9*”. To tolerate sequencing errors on corrected PacBio reads, parameters “*-motif TGCA -motifmis 1*” were used in related LTRharvest runs. To identify extra non-canonical LTR-RTs, no “*-motif*” parameter was specified for the maximum sensitivity.

Based on the annotation using the standard LTR library, the whole genome was categorized into four parts which are true positive (TP, LTR was identified), false negative (FN, LTR was not identified), false positive (FP, non-LTR was identified as LTR), and true negative (TN, non-LTR was not identified as LTR). Four metrics were used to evaluate the performance of LTR_retriever and its counterparts, which are sensitivity, specificity, accuracy, and precision defined as follows.

**Sensitivity = TP/(TP+FN)**

**Specificity = TN/(FP+TN)**

**Accuracy = (TP+TN)/(TP+TN+FP+FN)**

**Precision = TP/(TP+FP)**

Sensitivity, specificity, accuracy, and precision of each test were calculated using genomic sequence lengths by custom Perl scripts.

## DATA ACCESS

LTR_retriever is an open source software available in the GitHub repository (https://github.com/oushujun/LTR_retriever). Manually curated LTR libraries for rice and sacred lotus are available as supplementary files.

## FUNDING

This work was supported by the National Science Foundation [MCB-1121650 and IOS-1126998 to N.J.]; and the United States Department of Agriculture National Institute of Food and Agriculture and AgBioResearch at Michigan State University (Hatch grant MICL02120 to N.J.).

## ACKNOWLEDGMENTS

We thank Dr. Yi Liao (Institute of Genetics and Developmental Biology, Chinese Academy of Sciences) for valuable discussions. We thank Stefan Cerbin, Drs. Cornelius Barry, Rebecca Grumet, Steve van Nocker, and Wayne Loescher for critical reading of the manuscript.

